# *In vivo* fluorescence lifetime imaging captures metabolic changes in macrophages during wound responses in zebrafish

**DOI:** 10.1101/2020.06.16.153361

**Authors:** Veronika Miskolci, Kelsey E Tweed, Michael R Lasarev, Emily C Britt, Courtney E McDougal, Alex J Walsh, Jing Fan, John-Demian Sauer, Melissa C Skala, Anna Huttenlocher

**Author notes:** Co-corresponding Authors: Melissa Skala |, Anna Huttenlocher | (primary contact).

## Abstract

The effector functions of macrophages across the spectrum of activation states *in vitro* are linked to profound metabolic rewiring. However, the metabolism of macrophages remains poorly characterized *in vivo*. To assess changes in the intracellular metabolism of macrophages in their native inflammatory microenvironment, we employed two-photon fluorescence lifetime imaging microscopy (FLIM) of metabolic coenzymes NAD(P)H and FAD. We found that pro-inflammatory activation of macrophages *in vivo* was associated with a decrease in the optical redox ratio [NAD(P)H/(NAD(P)H+FAD)] relative to a pro-resolving population during both infected and sterile inflammation. FLIM also resolved temporal changes in the optical redox ratio and lifetime variables of NAD(P)H in macrophages over the course of sterile inflammation. Collectively, we show that non-invasive and label-free imaging of autofluorescent metabolic coenzymes is sensitive to dynamic changes in macrophage activation in interstitial tissues. This imaging-based approach has broad applications in immunometabolism by probing in real time the temporal and spatial metabolic regulation of immune cell function in a live organism.

**Significance:** Metabolic regulation of macrophage effector functions has recently emerged as a key concept in immune cell biology. Studies rely on *in vitro* and *ex vivo* approaches to study macrophage metabolism, however the high plasticity of these cells suggest that removal from their native microenvironment may induce changes in their intracellular metabolism. Here, we show that fluorescence lifetime imaging microscopy of metabolic coenzymes captures dynamic changes in the metabolic activity of macrophages while maintaining them in their endogenous microenvironment. This approach also resolves variations on a single-cell level, in contrast to bulk measurements provided by traditional biochemical assays, making it a potentially valuable tool in the field of immunometabolism.

## Introduction

Macrophages are innate immune cells from myeloid origin, distributed throughout most tissues of the body, and play key functions both in health and disease (1, 2). The heterogeneity and diversity in macrophage phenotypes and functions are well documented (3-5). Macrophages are commonly described in the literature as classically (M1) or alternatively (M2) activated, where M1 macrophages are associated with pro-killing functions such as eliminating pathogens and tumor cells, while M2 cells promote processes associated with wound healing and tumor progression. The simplistic nature of the M1/M2 classification, especially in context of *in vivo* biology, is controversial (6, 7), however it provides a framework amid the complexity of macrophage biology and remains widely used. Studies on macrophage activation recognized early on that M1 and M2 activation correlated with differential processing of some metabolites, most notably arginine (8). However, the importance of metabolic regulation of macrophage function was not fully appreciated until more recently, when it was recognized that some metabolic pathways are profoundly altered in classically activated macrophages (9, 10). This led to the emergence of metabolic reprogramming as a key hallmark of immune cell activation, that stresses that the metabolic state is not an outcome but a determinant of immune cell activation and function (11, 12). This is apparent from multiple metabolites functioning as signaling molecules outside of their traditional roles as intermediates in metabolic pathways (13). Evidence indicates that classically activated macrophages are glycolytic, while oxidative phosphorylation is the main fuel source during alternative activation (13, 14). Most studies in immunometabolism rely on using singular stimuli to activate macrophages, such as LPS or interleukin (IL)-4/IL-13, however macrophages face a complex mixture of stimuli within native tissues (15). This is further confounded by their plasticity, whereby macrophages readily adapt their responses to a changing microenvironment (16). This suggests that macrophage metabolism is best studied within the native microenvironment, however there are limited tools available to address this gap in understanding macrophage metabolism.

Nicotinamide adenine dinucleotide (NADH/NAD+) and flavin adenine dinucleotide (FADH_2_/FAD) are endogenous metabolic coenzymes, and serve as electron carriers in numerous metabolic pathways including glycolysis, the Krebs cycle, electron transport chain and oxidative phosphorylation (17, 18). These coenzymes are autofluorescent when reduced and oxidized, respectively, and allow for fluorescence lifetime imaging microscopy (FLIM) to quantify intracellular metabolism using intensity and lifetime measurements (17, 18). NADH and NADPH have overlapping spectral properties, and for accuracy NAD(P)H is used to reflect their combined signals (19). Fluorescence intensity can be used to determine the optical redox ratio that provides an assessment of the redox state of the cell (20). Multiple definitions of the optical redox ratio exist, but here we use NAD(P)H/(NAD(P)H+FAD), since an increase in the optical redox ratio intuitively corresponds with an increase in glycolysis, and it normalizes the values to be between 0 and 1 (21). Fluorescence lifetime measures the time a molecule spends in the excited state before decaying back to the ground state. Fluorescence lifetimes of NAD(P)H and FAD correlate with their enzyme-binding activities (17, 18), thereby reflecting changes in their cellular microenvironment. NADH and FAD exists in two forms, quenched and unquenched, resulting in short and long lifetimes. NAD(P)H has a short lifetime in the free state and a long lifetime in the protein-bound state (22). This is the converse for FAD, where FAD has a short lifetime in the bound state and a long lifetime in the free state. FLIM quantifies each of these lifetime components and the mean lifetime, defined by the weighted average of the short and long lifetimes. Fluorescence lifetime has several advantages over intensity measurements (23, 24). FLIM can provide additional biological information by distinguishing the protein-bound and free states, while NAD(P)H intensity is similar in both states. In addition, unlike intensity measurements, lifetime is independent of the cellular concentrations of the coenzymes. Nevertheless, intensity and lifetime in complement can quantify changes in cellular metabolism and have been used in a variety of applications (18). Importantly, FLIM is a label-free and non-invasive approach to detect metabolic changes *in situ* and can also resolve heterogeneity within a cell population based on the single cell-based imaging (25, 26).

Here, we explored whether we could use fluorescence lifetime imaging of NAD(P)H and FAD to assess changes in the metabolic activity of macrophages in a live animal, using larval zebrafish as our *in vivo* model. Zebrafish is well-suited to these studies given its high similarity to the human immune system (27) and genome (28), furthermore, the tail fin wounding is an established model of inflammation (29). Given the optical transparency at larval stage (29), this model is readily combined with fluorescence lifetime imaging to investigate the metabolic changes in macrophages over the course of an inflammatory response.

## Results

### *In vitro* validation of FLIM detects changes consistent with known metabolic profile

Our group and others have demonstrated that fluorescence lifetime measurements recapitulates the known metabolic profiles associated with *in vitro* macrophage activation in response to traditional stimuli such LPS/IFN-γ and IL-4/IL-13 (30-32). To further validate FLIM for detecting metabolic changes in a whole organism, first we tested whether we could recapitulate previously reported findings on macrophage activation *in vitro* in response to an intracellular pathogen. Gillmaier *et al* used ^13^C-isotopologue profiling to trace carbon metabolism during infection of primary mouse macrophages with the intracellular pathogen, *Listeria monocytogenes* (*Lm*), and found that infection is associated with increased glycolytic activity in the host cells (33). In a similar fashion, we infected mouse bone marrow derived macrophages (BMDM) with *L. monocytogenes* and performed fluorescence lifetime imaging of NAD(P)H and FAD on live cells at 5-6 hours post infection (hpi). Intracellular growth of *L. monocytogenes* peaks by 5 hpi and remains at a plateau, and by 8 hpi a fraction of macrophages undergo cell death (34). Intracellular infection was monitored by mCherry labeling of *L. monocytogenes*. The optical redox ratio is a measure of the oxidation-reduction state of the cells, and increased glycolytic rate generates NADH leading to an increase in NADH levels, thereby an increase in the redox ratio [as defined here NAD(P)H/(NAD(P)H + FAD)] (17, 18). As expected, we detected a slight but significant increase in the optical redox ratio of infected macrophages compared to uninfected control (Figure 1A, B; the mCherry signal was used to subtract the bacterial metabolic data (see figure legend and methods) to exclude from macrophage metabolism). This change in the optical redox ratio is consistent with an increase in glycolytic activity of the infected host cells measured by ^13^C-isotopologue profiling. *L monocytogenes* infection of macrophages resulted in the increase of the mean lifetime (τ_m_) of NAD(P)H, but not FAD (Figure 1C, D). We also observed alterations in the individual lifetime components that were associated mostly with NAD(P)H (Figure S1A-F).

**Figure 1.**
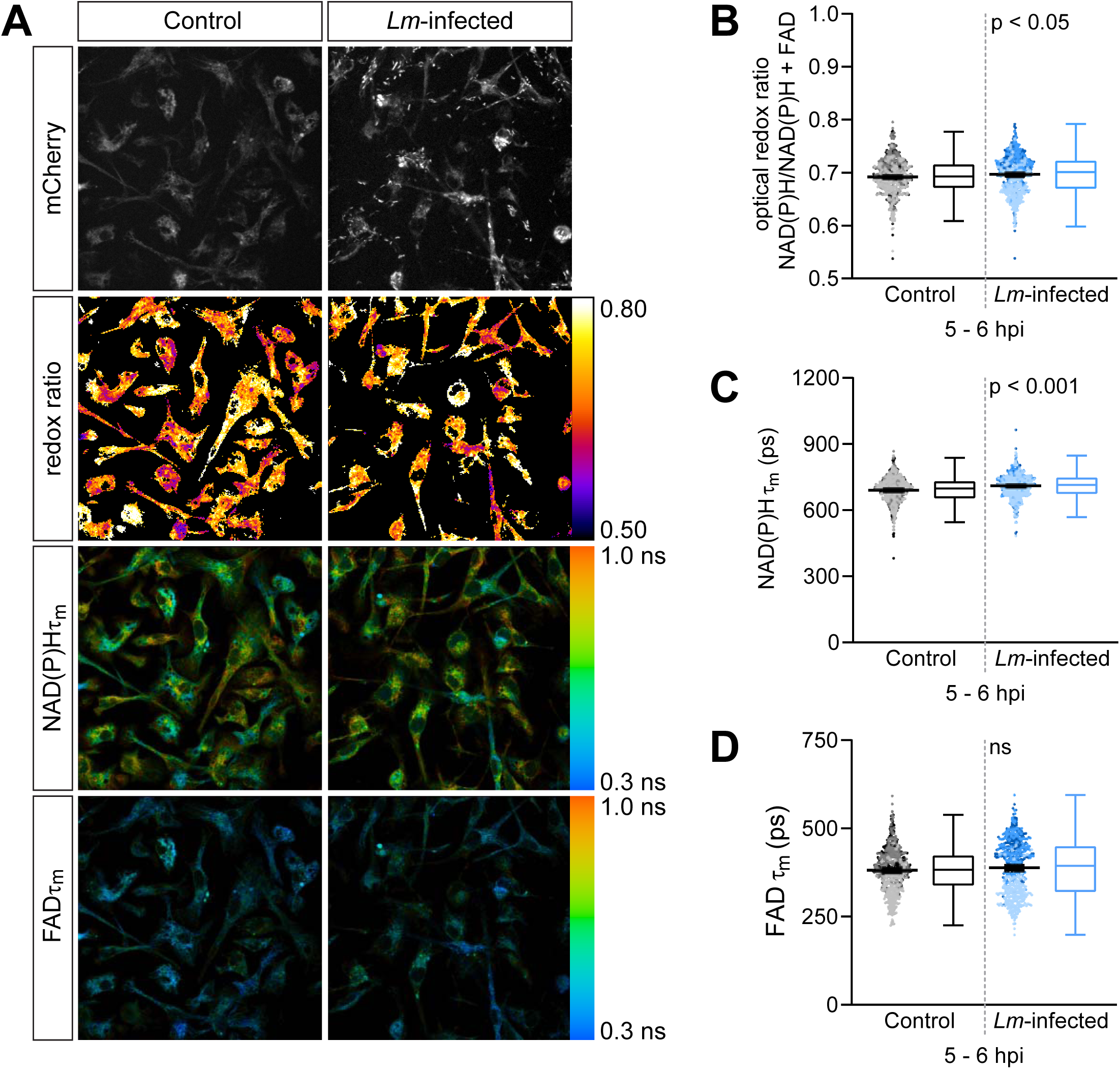
Fluorescence lifetime imaging of NAD(P)H and FAD detects predicted changes in the optical redox ratio in the context of *in vitro* infection of primary macrophages. Mouse bone marrow derived macrophages were infected with mCherry-labeled *Listeria monocytogenes (Lm)* at MOI 2, and FLIM of NAD(P)H and FAD was performed on live cells at 5-6 hours post infection (hpi). A) Representative images of mCherry (to show presence of bacteria), optical redox ratio, and NAD(P)H and FAD mean lifetimes (τ_m_) are shown for uninfected control or *Lm*-infected macrophages; scale bar = 50 µm. Quantitative analysis of B) optical redox ratio, C) NAD(P)H and D) FAD mean lifetimes (τ_m_ = α_1_τ_1_ + α_2_τ_2_) are shown; quantitative analysis of other associated lifetime endpoints (α_1_, τ_1_, τ_2_) are included in Figure 1 supplement. The diffuse cytoplasmic fluorescence in the mCherry images is likely due to FAD autofluorescence (44). mCherry expression in bacteria was used to subtract from lifetime images in order to exclude metabolic changes in the pathogen from that of macrophages. Results from 3 independent repeats are shown; sample size for each repeat is included in Figure 1 supplement. Each data point represent a macrophage and the data for each condition is displayed by a composite dotplot and boxplot; each repeat is displayed by a different color in the dotplot, showing mean with 95% CI; boxplots show median (central line), first and third quartiles (lower and upper lines), and the Tukey method was employed to create the whiskers (the farthest data points that are no further than 1.5 times the interquartile range); data points beyond whiskers (refer to dotplot) are considered outliers. Statistical comparison was performed by general linear model; ns = not significant.

### *In vivo* validation of FLIM with metabolic inhibitor treatment produces predicted changes in the optical redox ratio

To further validate *in vivo* detection of metabolic changes in macrophages, next we carried out metabolic inhibitor treatment in our *in vivo* system to monitor for predicted changes in the optical redox ratio of macrophages responding to a tail wound in larval zebrafish. To be able to segment autofluorescence signal associated with macrophages from the whole tissue, we used a transgenic reporter line where macrophages are identified by fluorescent protein expression. We have empirically tested the compatibility of several fluorescent proteins with the spectral properties of NAD(P)H and FAD (data not shown). We found that GFP is suitable to image in conjunction with NAD(P)H, but it excludes the acquisition of FAD as they have overlapping spectra. However, we found mCherry to be compatible for imaging with NAD(P)H and FAD. We also optimized imaging on live larvae. Serial acquisition of NAD(P)H and FAD was not suitable for imaging motile cells, such as macrophages, in live larvae (35). To accommodate cell movement during image acquisition in live larvae, we employed the wavelength mixing approach that allows for simultaneous acquisition in three different channels (36). We performed simple tail fin transection on transgenic larvae (*Tg(mpeg1:mCherry-CAAX)* that labels macrophages with mCherry), and performed fluorescence lifetime imaging of NAD(P)H and FAD at the wound region (Figure 2A) at 3-6 hours post tail transection (hptt) in the absence or presence of glycolysis inhibitor 2-deoxy-d-glucose (2-DG). 2-DG is a glucose analog and acts as a competitive inhibitor of glycolysis at the step of phosphorylation of glucose by hexokinase (37). As inhibiting glycolysis reduces NADH levels (17, 18), we expected the optical redox ratio to decrease in macrophages of treated larvae compared to untreated control. Indeed, the optical redox ratio was significantly lower in macrophages in the 2-DG-treated larvae (Figure 2B, C). This change was driven by a decrease in NAD(P)H intensity in treated larvae, while FAD intensity remained similar to control levels (data not shown). NAD(P)H τ_m_ significantly decreased in macrophages of treated larvae, while the change for FAD τ_m_ was marginal (Figure 2D, E). We also observed significant changes in some of the individual lifetime components of NAD(P)H and FAD (Figure S2A-F). These effects on NAD(P)H and FAD lifetime endpoints were similar to the effects observed during 2-DG treatment of activated T cells (21). In sum, upon inhibition of glycolysis with 2-DG in our *in vivo* tail wound model, we observed the expected changes in the optical redox ratio in macrophages responding to sterile wounds, supporting the utility of this approach to detect changes in macrophage metabolism in interstitial tissues of live animals.

**Figure 2.**
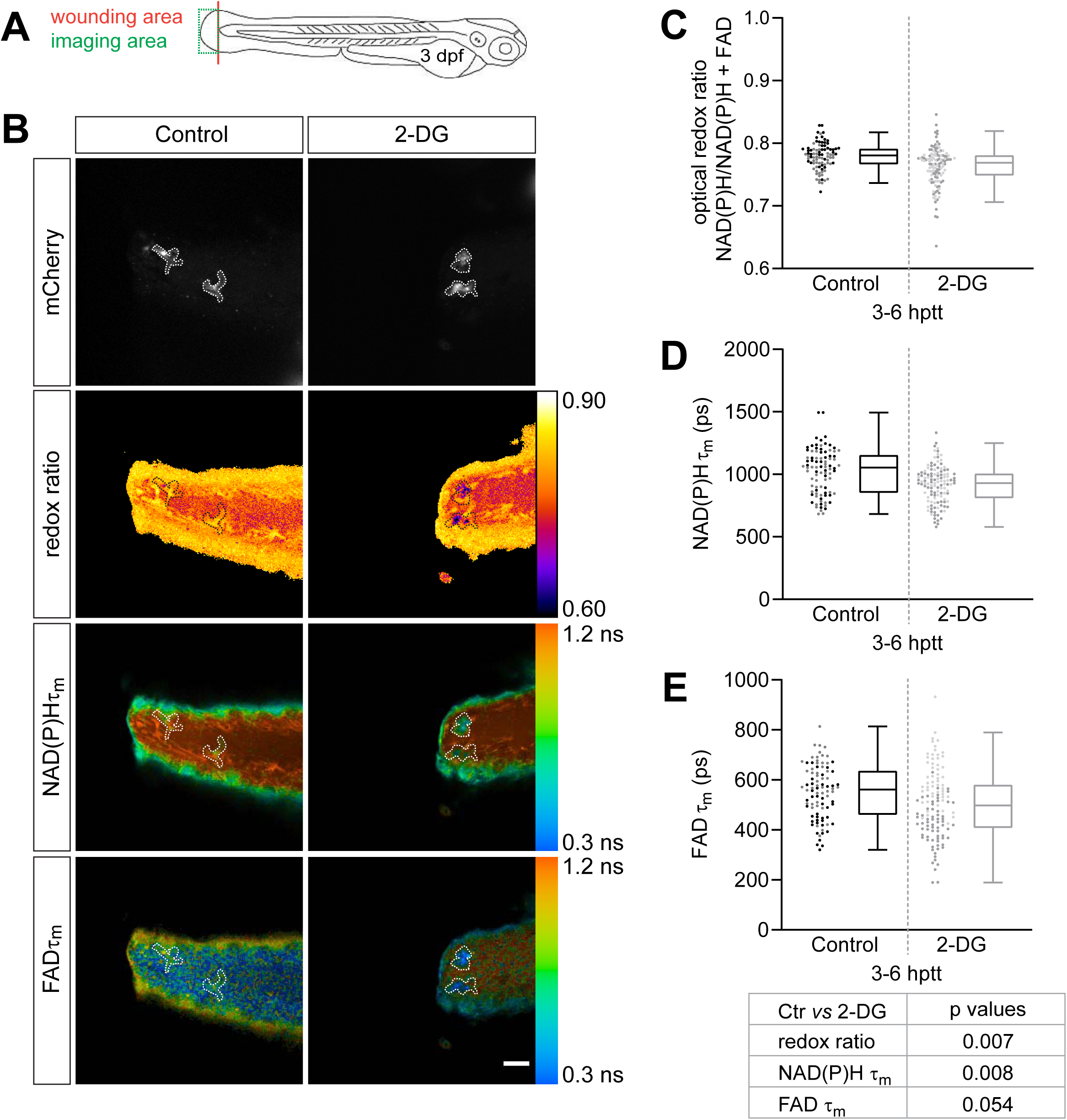
Inhibition of glycolysis produces predicted changes in the optical redox ratio of macrophages at the simple tail wound. Tail fin transection distal to the notochord was performed using transgenic zebrafish larvae (*Tg(mpeg1:mCherry-CAAX)* that labels macrophages in the plasma membrane with mCherry) at 3 days post fertilization (dpf), and FLIM of NAD(P)H and FAD was performed on live larvae at 3-6 hours post tail transection (hptt) that were either untreated (control) or treated with 5 mM 2-DG (glycolysis inhibitor). A) Schematic showing area where wounding (red line) and imaging (green box) was performed. B) Representative images of mCherry (to show macrophages), optical redox ratio, and NAD(P)H and FAD mean lifetimes (τ_m_) are shown for control or treated tail wounds; macrophages in mCherry channel were outlined with dashed lines and the area was overlaid in the optical redox ratio and lifetime images to show corresponding location; scale bar = 50 µm. Quantitative analysis of C) optical redox ratio, D) NAD(P)H and E) FAD mean lifetimes (τ_m_ = α_1_τ_1_ + α_2_τ_2_) are shown; quantitative analysis of other associated lifetime endpoints (α_1_, τ_1_, τ_2_) are included in Figure 2 supplement. Results from 2 biological repeats are shown; sample size for each repeat is included in Figure 2 supplement. Each data point represent a macrophage and the data for each condition is displayed by a composite dotplot and boxplot; each repeat is displayed by a different color in the dotplot; boxplots show median (central line), first and third quartiles (lower and upper lines), and the Tukey method was employed to create the whiskers (the farthest data points that are no further than 1.5 times the interquartile range); data points beyond whiskers (refer to dotplot) are considered outliers. Statistical comparison was performed using a general linear model with cluster-robust standard errors to account for multiple macrophages measured per larvae, thereby statistical conclusions are shown in a table below the graphs. The optical redox ratio and τ_m_ were log-transformed prior to analysis. Estimated means with 95% CI are included in Figure 2 supplement.

### Fluorescence lifetime imaging detects metabolic changes in macrophage subsets at infected tail wounds

To begin to address whether fluorescence lifetime imaging could distinguish different macrophage populations in a whole organism, we decided to use our recently developed zebrafish infected tail wound model (38). This allows us to induce the recruitment of differentially polarized macrophages under physiological conditions. The infected tail wound model combines the simple tail fin transection with *L. monocytogenes* infection, and provokes an extensive and sustained infiltration of macrophages, that is characterized by a large portion of macrophages expressing high levels of TNFα (38). TNFα is a well-established marker of the M1-like pro-killing phenotype of macrophages across several species, including zebrafish (3, 39). This inflammatory response is unlike what we observe following the simple tail fin transection, where most macrophages are TNFα− throughout the course of the wound response (38). Due to the lack of TNFα expression, the macrophages at the simple tail wound are presumed to represent differentially polarized macrophages, most likely the M2-like (39). Based on the recruitment of differentially polarized macrophage populations by these tail wound models, we hypothesized that we would detect differences in the metabolic activity of macrophages between the simple and *L. monocytogenes*-infected tail fin wounds. We performed tail fin transection in the absence or presence of *L. monocytogenes* on double transgenic (*Tg(tnf:GFP) x Tg(mpeg1:mCherry-CAAX)*) larvae, and performed lifetime imaging of NAD(P)H at the wound region on live larvae at 48 hours post wound (hpw). In this set of experiments we performed lifetime imaging in conjunction with the TNFα reporter line (*tnf:GFP*) in order to monitor and group macrophages by TNFα expression during data analysis. The TNFα reporter line relies on GFP expression to report transcriptional activity of *tnfa* (40), which precludes acquisition of FAD measurements. As a result, in this experiment we were not able to monitor changes in the intracellular optical redox ratio. Macrophages at the wound region were identified based on plasma membrane-localized mCherry expression as above. Since the adaptive immune system has not developed yet at this larval stage, the polarized activation of macrophages in these experiments does not reflect the involvement of T cells (27). As previously reported (38), the infected tail wound recruited significantly more macrophages compared to the uninfected control (simple tail wound). While both tail wounds recruited a macrophage population with mixed levels of TNFα expression, macrophages at the infected tail wound were significantly more M1-like as most cells had high TNFα expression, while the majority were TNFα− at the uninfected control (Figure 3A, Figure S3C). We detected a significant reduction in the mean lifetime of NAD(P)H for TNFα+ relative to TNFα− macrophages from either the uninfected control or *Lm*-infected tail wound (Figure 3B). NAD(P)H τ_m_ was also significantly reduced in macrophages from *Lm*-infected tail wounds relative to uninfected control when comparing either the TNFα− or TNFα+ groups (Figure 3B). The trends for the differences in the individual lifetime components of NAD(P)H were similar to that observed for τ_m_ (Figure S3A, B), while we did not detect any significant changes in the fractional component of free NAD(P)H (α_1_) in any of the comparisons (Figure 3C). Next, we repeated the same set of experiments but without the TNFα reporter line, in order to acquire FAD lifetime measurements and monitor changes in the optical redox ratio. We detected a significant reduction in the mean lifetime and other individual lifetime components of NAD(P)H in macrophages at the wound of the *Lm*-infected larvae (Figure 4C, Figure S4B, C), consistent with the measurements above (Figure 3B, Figure S3A, B); under these conditions, we found that NAD(P)H α_1_ significantly increased in macrophages at the *Lm*-infected wound (Figure S4A). The presence of infection at the tail wound did not induce any significant changes in FAD lifetime endpoints (Figure 4D, Figure S4D-F). Interestingly, the optical redox ratio was significantly reduced in macrophages in the highly inflammatory *Lm*-infected wound as compared to the uninfected control wound (Figure 4A, B). This result was unexpected considering the observed increase of the redox ratio in the context of the *in vitro* infection of BMDM with *L. monocytogenes* (Figure 1B). We reasoned this result may be influenced by the presence of an intracellular pathogen in macrophages at the *Lm*-infected tail wound, and not solely due to a more inflammatory macrophage phenotype. To test this, we proceeded to measure changes in the cellular metabolism of macrophages in context of a sterile tail wound that induces the recruitment of TNFα+ macrophages.

**Figure 3.**
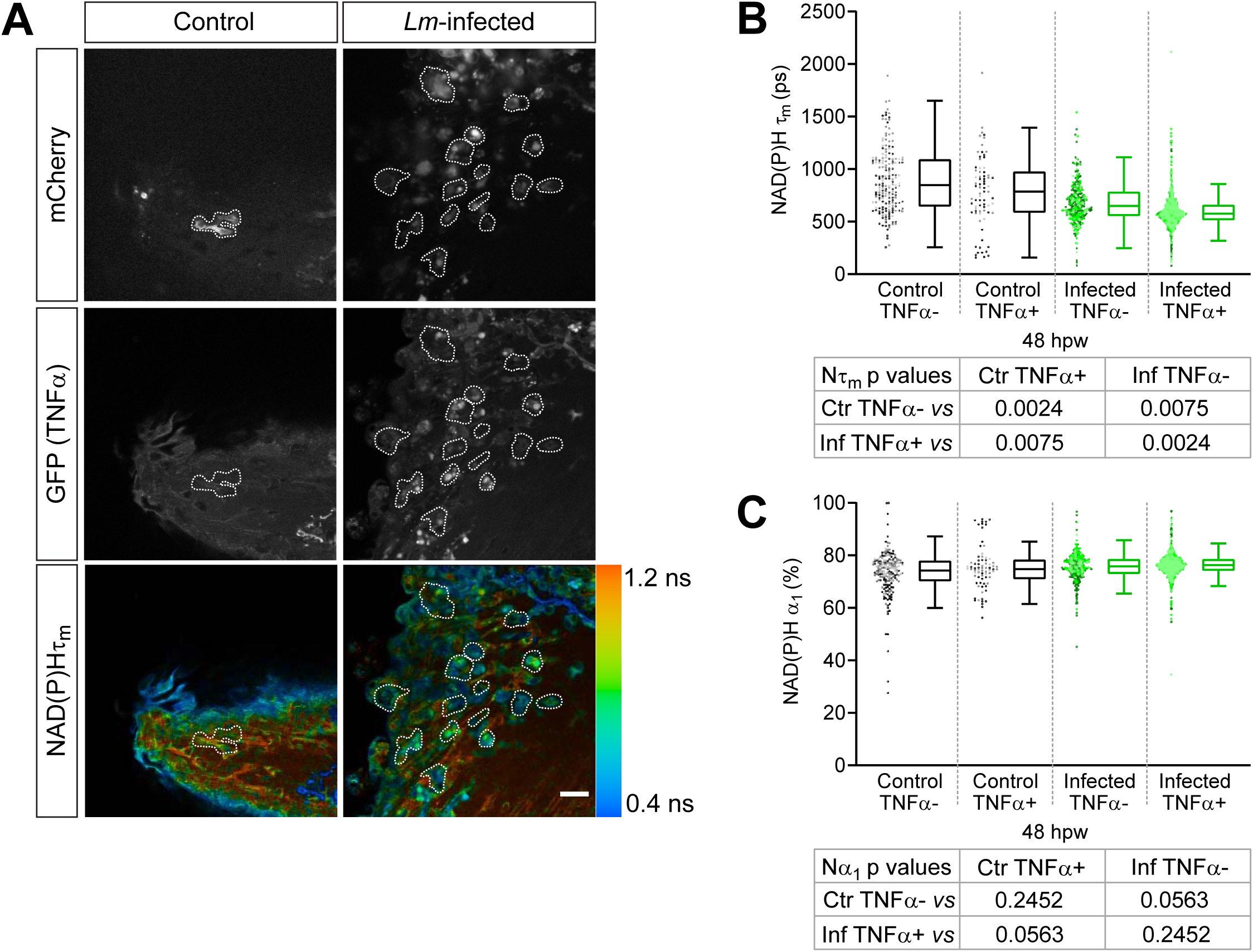
Fluorescence lifetime imaging of NAD(P)H detects metabolic changes in TNFα− and TNFα+ macrophages at the infected tail wound. Tail fin transection distal to the notochord was performed using double transgenic zebrafish larvae (*Tg(tnf:GFP) x Tg(mpeg1:mCherry-CAAX)*, a TNFα reporter line in combination with a line that labels macrophages in the plasma membrane with mCherry) at 3 days post fertilization (dpf) in the absence or presence of *Listeria monocytogenes (Lm)*. FLIM of NAD(P)H was performed on live larvae at 48 hours post wound (hpw) (Figure 2A). A) Representative images of mCherry expression to show macrophages, GFP to show TNFα expression and NAD(P)H mean lifetime (τ_m_) are shown for control or infected tail wounds; macrophages in mCherry channel were outlined with dashed lines and the area was overlaid in GFP and lifetime images to show corresponding location; in the infected condition only few macrophages are outlined as examples; scale bar = 50 µm. Quantitative analysis of B) NAD(P)H mean lifetime (τ_m_ = α_1_τ_1_ + α_2_τ_2_) and C) alpha1 (α_1_), fractional component of free NAD(P)H are shown; quantitative analysis of other associated lifetime endpoints (τ_1_, τ_2_) are included in Figure 3 supplement. Results from 3 biological repeats are shown; sample size for each repeat is included in Figure 3 supplement. Each data point represent a macrophage and the data for each condition is displayed by a composite dotplot and boxplot; each repeat is displayed by a different color in the dotplot; boxplots show median (central line), first and third quartiles (lower and upper lines), and the Tukey method was employed to create the whiskers (the farthest data points that are no further than 1.5 times the interquartile range); data points beyond whiskers (refer to dotplot) are considered outliers. Statistical comparison was performed using a general linear model with cluster-robust standard errors to account for multiple macrophages measured per larvae, thereby statistical conclusions are shown in a table below the graphs. The lifetime endpoints were log-transformed prior to analysis. Interaction between treatment and GFP expression was included to analyze whether either factor modified the effect of the other; no interaction was found. Estimated means with 95% CI are included in Figure 3 supplement.

**Figure 4.**
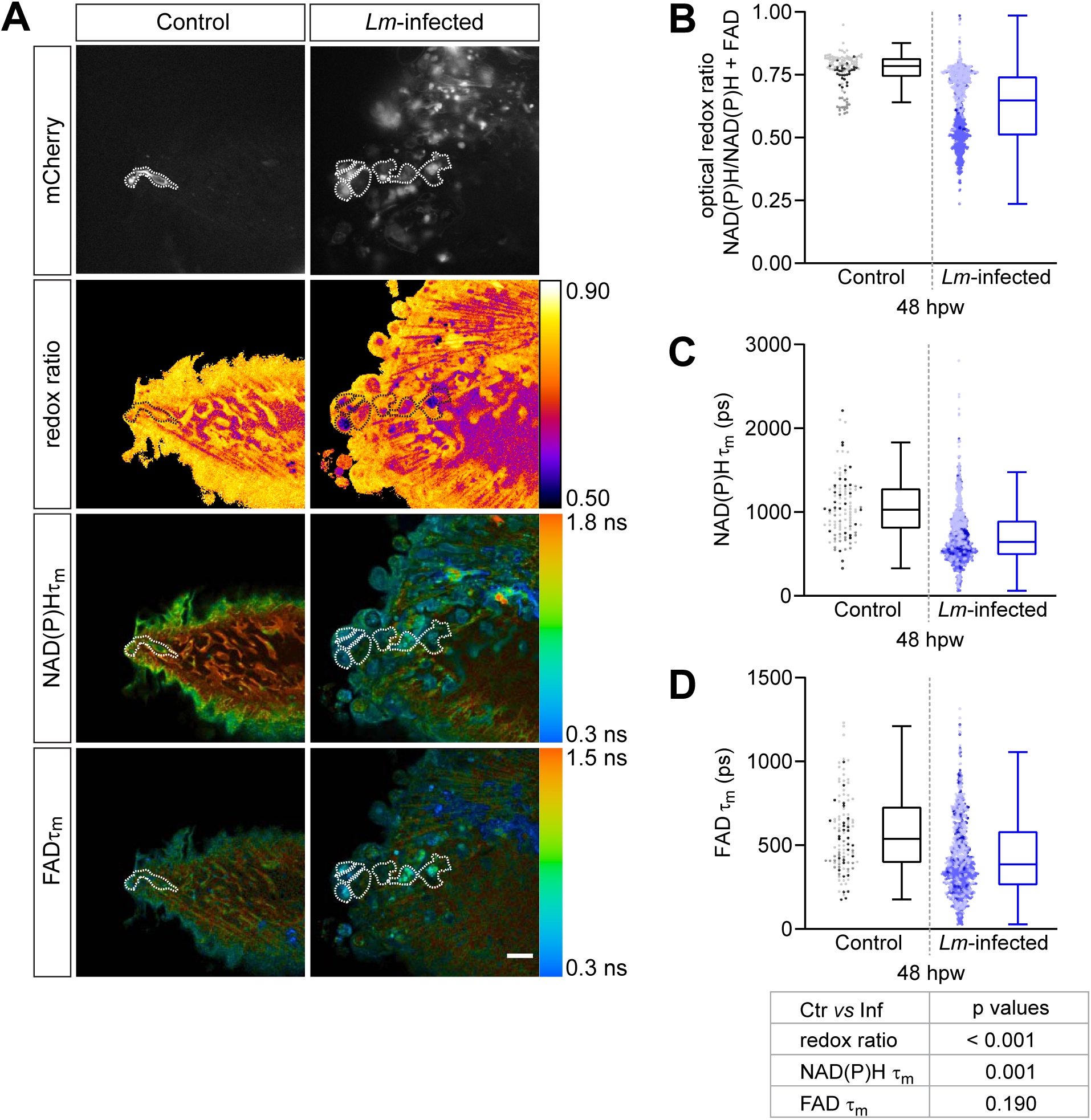
Fluorescence lifetime imaging of NAD(P)H and FAD detects metabolic changes in macrophages at the infected tail wound. Tail fin transection distal to the notochord was performed using transgenic zebrafish larvae (*Tg(mpeg1:mCherry-CAAX)* that labels macrophages in the plasma membrane with mCherry) at 3 days post fertilization (dpf) in the absence or presence of *Listeria monocytogenes (Lm)*. FLIM of NAD(P)H and FAD was performed on live larvae at 48 hours post wound (hpw) (Figure 2A). A) Representative images of mCherry expression to show macrophages, optical redox ratio, and NAD(P)H and FAD mean lifetimes (τ_m_) are shown for control or infected tail wounds; macrophages were outlined with dashed lines and the area was overlaid in the optical redox ratio and lifetime images to show corresponding area; in the infected condition only few macrophages are outlined as examples; scale bar = 50 µm. Quantitative analysis of B) optical redox ratio, C) NAD(P)H and D) FAD mean lifetimes (τ_m_ = α_1_τ_1_ + α_2_τ_2_) are shown; quantitative analysis of other associated lifetime endpoints (α_1_, τ_1_, τ_2_) are included in Figure 4 supplement. Results from 3 biological repeats are shown; sample size for each repeat is included in Figure 4 supplement. Each data point represent a macrophage and the data for each condition is displayed by a composite dotplot and boxplot; each repeat is displayed by a different color in the dotplot; boxplots show median (central line), first and third quartiles (lower and upper lines), and the Tukey method was employed to create the whiskers (the farthest data points that are no further than 1.5 times the interquartile range); data points beyond whiskers (refer to dotplot) are considered outliers. Statistical comparison was performed using a general linear model with cluster-robust standard errors to account for multiple macrophages measured per larvae, thereby statistical conclusions are shown in a table below the graphs. Log transformation was applied to τ_m_ prior to analysis. Estimated means with 95% CI are included in Figure 4 supplement.

### Fluorescence lifetime imaging of NAD(P)H and FAD resolves changes in the metabolic activity of macrophages over the course of a sterile inflammatory response

To measure metabolic activity of macrophages during sterile inflammation, we used our recently developed zebrafish thermal injury tail wound model (38). This injury is produced by briefly burning the tail fin tissue distal to the notochord using a surgical cautery wire. Similarly to the infected wound, the burn wound also elicits the recruitment of a more M1-like macrophage population compared to the simple transection, where a large percentage of macrophages are TNFα+ (38). Unlike at the infected wound where expression of TNFα persists, TNFα+ macrophages peak at 24 hpw and resolve thereafter, with most macrophages at the wound being TNFα− by 72 hpw following thermal injury (38). We performed lifetime imaging of NAD(P)H and FAD at these two time points to compare the metabolic activity of macrophages in response to simple transection and thermal injury. Since macrophages are mostly TNFα− throughout the course of the wound response following a simple transection, we speculated that the metabolic activity of macrophages would be different at 24 hpw, but not at 72 hpw, at the two wounds. We performed tail fin transection or burn wound distal to the notochord on transgenic (*Tg(mpeg1:mCherry-CAAX)*) larvae, and performed lifetime imaging at the wound region on live larvae at 24 and 72 hpw (Figure 5A). As expected, we observed significant differences in the metabolic activity of macrophages between the wounds at 24 hpw, but the cellular metabolism was similar at 72 hpw. Importantly, at 24 hpw the optical redox ratio was lower in macrophages at the burn wound relative to macrophages at the simple transection (Figure 5B), similar to what was observed at the infected wound (Figure 4B). The trends for the differences in the mean lifetime and individual lifetime components of NAD(P)H and FAD between the burn wound and the simple transection was also similar to what we observed between the infected wound and the simple transection (Figure 5C, D, Figure S5A-F). As expected, the optical redox in macrophages was not different between the simple transection and burn wound at 72 hpw (Figure 5B). Macrophages are mostly TNFα− at both wounds by 72 hpw (38), suggesting that the macrophage populations present at these wounds have similar activation states and thereby likely to have similar metabolic activity. The mean lifetime of NAD(P)H was significantly lower in macrophages at the burn wound relative to the simple transection at 72 hpw, while it was similar for FAD (Figure 5C, D); most of individual lifetime components of NAD(P)H and FAD were also similar between the two wounds at 72 hpw (Figure S5A-F). Generally, most of the changes in lifetime endpoints were associated with NAD(P)H, and they were consistent across the different wound models (Table 1). Furthermore, we also detected temporal changes in the metabolic activity of macrophages during the wound responses. The optical redox ratio of macrophages increased over time in response to the simple transection, as well as to thermal injury (Figure 5B). This would be expected as the macrophage population at the burn wound becomes more like macrophages at simple transection over time, based on TNFα expression (38). In line with this, the trends for the changes in the lifetime endpoints over time were also similar at the two wounds. NAD(P)H τ_m_ increased and FAD τ_m_ decreased over time at both wounds (Figure 5C, D); the trends for changes in the individual lifetime components of NAD(P)H and FAD over time were also similar at the two wounds (Figure S5A-F).

**Table 1.** Summary of changes in optical redox ratio and NAD(P)H lifetime endpoints.

|  | *in vitro*<br>*Lm*-infection<sup>1</sup> | 2-DG<sup>2</sup> | *Lm*-infected wound<sup>2</sup> |  | *Lm*-infected<br>wound<sup>2</sup> | Burn wound<sup>2</sup> |  |
| --- | --- | --- | --- | --- | --- | --- | --- |
|  |  |  | TNFα- | TNFα+ |  | 24h | 72h |
| Redox ratio | ↑ | ↓ | na | na | ↓ | ↓ | nd |
| NAD(P)H τ<sub>m</sub> | ↑ | ↓ | ↓ | ↓ | ↓ | ↓ | ↓ |
| NAD(P)H τ<sub>1</sub> | ↑ | ↓ | ↓ | ↓ | ↓ | ↓ | nd |
| NAD(P)H τ<sub>2</sub> | ↑ | ↓ | ↓ | ↓ | ↓ | ↓ | nd |
| NAD(P)H α<sub>1</sub> | ↓ | ↑ | ↑ | ↑ | ↑ | ↑ | ↑ |

Changes in treated samples are shown relative to respective controls. Control = 1-uninfected BMDM, 2-tail fin transection wound (uninfected); na = not applicable; nd = not different.

**Figure 5.**
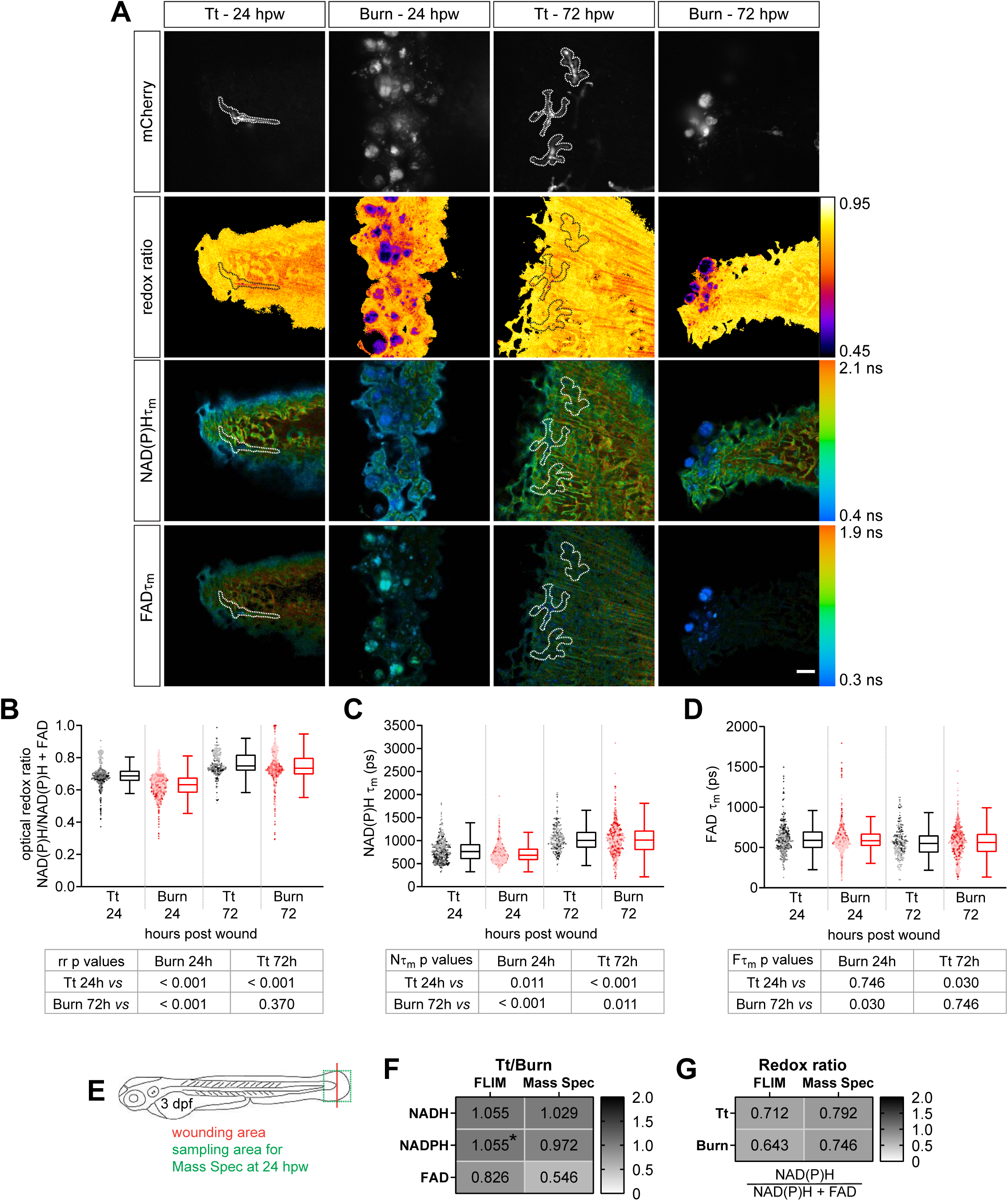
Fluorescence lifetime imaging of NAD(P)H and FAD detects temporal changes in the metabolic activity of macrophages at sterile tail wounds. Tail fin transection (Tt) or thermal injury (Burn) distal to the notochord was performed using transgenic zebrafish larvae (*Tg(mpeg1:mCherry-CAAX)* that labels macrophages in the plasma membrane with mCherry) at 3 days post fertilization (dpf). FLIM of NAD(P)H and FAD was performed on live larvae at 24 and 72 hours post wound (hpw) (Figure 2A). A) Representative images of mCherry expression to show macrophages, optical redox ratio, and NAD(P)H and FAD mean lifetimes (τ_m_) are shown for Tt or Burn wounds; macrophages were outlined with dashed lines and the area was overlaid in the optical redox ratio and lifetime images to show corresponding location of macrophages; scale bar = 50 µm. Quantitative analysis of B) optical redox ratio, C) NAD(P)H and D) FAD mean lifetimes (τ_m_ = α_1_τ_1_ + α_2_τ_2_) are shown; quantitative analysis of other associated lifetime endpoints (α_1_, τ_1_, τ_2_) are included in Figure 5 supplement. Results from 3 biological repeats are shown; sample size for each repeat is included in Figure 5 supplement. Each data point represent a macrophage and the data for each condition is displayed by a composite dotplot and boxplot; each repeat is displayed by a different color in the dotplot; boxplots show median (central line), first and third quartiles (lower and upper lines), and the Tukey method was employed to create the whiskers (the farthest data points that are no further than 1.5 times the interquartile range); data points beyond whiskers (refer to dotplot) are considered outliers. Statistical comparison was performed using a general linear model with cluster-robust standard errors to account for multiple macrophages measured per larvae, thereby statistical conclusions are shown in a table below the graphs. Log transformation was applied to τ_m_ prior to analysis. Interaction between treatment and time was included to analyze whether either factor modified the effect of the other; Strong interaction was detected for the optical redox ratio. E) Tail fin tissue was collected distal to the caudal vein/artery loop (green box) 24 h following either tail fin transection or thermal injury distal to the notochord (red line) for mass spec analysis of small metabolites to compare the global trend of changes in redox metabolites with that measured by FLIM; metabolomics data shown in F and G are from 4 biological repeats. F) metabolite abundance measured by either fluorescence intensity (FLIM) or mass spec in transection sample was normalized by that in burn or G) was used to calculate the redox ratio in transection (Tt) or burn samples. We included NADPH abundance in the redox ratio calculated using Mass Spec measurements. *NADPH and NADH intensities were not collected separately by FLIM as their fluorescence spectra overlap, thereby measured collectively. Estimated means with 95% CI are included in Figure 5 supplement.

To further substantiate the metabolic changes observed in macrophages at the sterile tail wounds, we tested whether we could recapitulate our FLIM measurements with a different method. Fluorescence intensities of NAD(P)H and FAD detected by FLIM is a measure of their relative intracellular abundance. Similarly, we analyzed the abundance of NADH, NADPH and FAD by targeted metabolomics (41). We performed tail fin transection or burn wound distal to the notochord on wild-type zebrafish larvae and collected the tail fin tissue distal to the caudal vein/artery loop (to remain close to the wound microenvironment) 24 hours following injury for targeted metabolomics (Figure 5E). The technical limitation here is that we analyzed the abundance of these small metabolites in the whole tail fin tissue, not macrophages alone, because it is difficult to collect sufficient numbers of macrophages from such a small region to reach the detection limit of the mass spectrometer. We calculated the relative abundances of NAD(P)H and FAD in burn wound compared to transection, and found the trends measured by both FLIM and mass spectrometry are consistent, i.e. the NAD(P)H level are similar in both wound models, while FAD level is lower in transection (Figure 5F). Additionally, we compared the redox ratio measured in each wound model by mass spectrometry and FLIM (Figure 5G). The two methods gave similar results, and both showed the trend that the redox ratio is higher in transection. The differences in redox ratio values measured by these two methods is likely due to the fact that our metabolomics method measures the whole tail fin tissue of wound microenvironment, while FLIM measures the wound-associated macrophages specifically.

## Discussion

Immunometabolism has become a fast-growing and exciting field based on the realization that metabolism plays a profound regulatory role in immune cell activation (11, 12). New therapies are focused on metabolic targets in immune cells, such as macrophages, which play a key role in autoimmunity and the progression of human diseases such as arthritis, atherosclerosis and cancer (12, 42, 43). *In vitro* analyses have formed the foundation of immunometabolism and have provided fundamental insights into the metabolic regulation of immune cell biology. However *in vitro* studies fail to reflect the complexities associated with *in vivo* environments, including the input from mixed signals and interactions with other cells (15). An understanding of *in vivo* behavior has been hampered by the lack of tools available for the *in vivo* assessment of functional metabolic changes. As a result, immunometabolism *in vivo* remains poorly characterized. Lifetime imaging of the endogenous fluorescence of metabolic coenzymes is an attractive approach because it allows for the quantitative analysis of metabolic changes on a single-cell level, while maintaining cells in their native microenvironment. Studies on the metabolic profiles of macrophages *in vivo* using FLIM have been limited, with one study demonstrating that macrophages have distinguishable lifetime signatures from tumor cells at the tumor microenvironment (44). Here, we took advantage of complex *in vivo* wound models we recently developed that each induce a characteristic macrophage inflammatory response (38). This is, to our knowledge, is the first study to examine the potential of fluorescence lifetime imaging to distinguish macrophages with differential activation states within interstitial tissues in live animals.

Our findings suggest that macrophage metabolism *in vitro* and *in vivo* may differ. Macrophages infected by *L. monocytogenes in vitro* exhibited increased optical redox ratio relative to uninfected macrophages (Figure 1B), however we found that infection of live zebrafish reduced the optical redox ratio in macrophages (Figure 4B). Surprisingly, we found that pro-inflammatory macrophages even at sterile inflammatory sites were also associated with reduced optical redox ratios (Figure 5B), suggesting that a pro-inflammatory macrophage population *in vivo* has more oxidative metabolic state in general. We characterized macrophage populations in live zebrafish larvae using the TNFα reporter (40), and categorized cells with high TNFα expression as M1-like. Currently, there are no live reporters to mark the M2-like macrophage population in larval zebrafish, and the lack of TNFα expression does not confirm that. Nevertheless, we consider TNFα− and TNFα+ macrophages as different populations of cells. In light of the known *in vitro* metabolic profiles of macrophages, we expected a macrophage population with large number TNFα+ cells to exhibit a more glycolytic state and thereby have higher redox ratio relative to a mostly TNFα− macrophage population. One caveat of intensity-based measurements is that other fluorophores, such as elastin and lipofuscin, could contribute to the intensity signals for the redox ratio and be a source of error (24). The source and role of the observed oxidative metabolism in macrophages in the context of infection and sterile inflammation *in vivo* require further analysis. Our initial analysis suggests that mitochondrial reactive oxygen species (mROS) contributes to the observed NAD(P)H and FAD lifetime profiles during sterile inflammation without affecting the redox state of the cell (data not shown), and it will be interesting to further explore the role of mROS in macrophage activation and function *in vivo*.

Interestingly, we found that TNFα expression in macrophages at the infected tail wound was associated with a graded effect on NAD(P)H lifetime endpoints that was on a spectrum (Figure 3B, Figures S3A, B), where TNFα− macrophages from uninfected wound (control) are on one end of the spectrum, while TNFα+ cells from the infected wound are on the opposite end. As we move on this spectrum, we see a graded change in the lifetime endpoints in the same direction from one end to the other end, reminiscent of the concept that macrophage activation *in vivo* occurs on a spectrum as opposed to a more strict M1 or M2 (3). These results suggest that FLIM is sensitive to variations in macrophage populations across different levels of activation. Furthermore, we found that fluorescence lifetime imaging is also able to resolve time-related changes in macrophage metabolism. We observed that the redox ratio in macrophages increased over time, both at the simple tail wound and the burn wound (Figure 5B). The observation that changes in metabolic activity of macrophages at the burn wound resembles the changes at the simple tail wound is expected, as the macrophage population at the burn wound becomes similar to that at the simple wound over time (38). Previously we described that the macrophage population at the burn wound is mostly TNFα+, however over time the macrophage population becomes mostly TNFα−, similar to the simple wound, at this switch in activation phenotype coincided with a recovery in wound healing (38). The observed increase in the optical redox ratio over time is interesting. Macrophages polarize towards a pro-healing M2-like state during wound healing, based on which we expected macrophages to have a more oxidative metabolism (11, 14) and thereby display a reduction in the optical redox ratio. However, it has been demonstrated that M2-like macrophages are more motile compared to M1-like cells (45) and glycolytic reprogramming has been shown to be important for macrophage migration (46). These reports suggest that the observed increase in the optical redox ratio, reflecting an increase in glycolytic activity, may be supporting the more motile nature of pro-healing M2-like cells.

With the emergence of immunometabolism it is now recognized that metabolic reprogramming underlies macrophage activation and function. Differential activation of macrophages plays a central role in host health and disease progression, underscoring the importance of studying macrophage metabolism *in vivo*. We have shown that fluorescence lifetime imaging of NAD(P)H and FAD is able to resolve metabolic changes in macrophages with distinct activation states *in situ* in a live organism, suggesting that FLIM can be a valuable imaging-based tool to study the metabolic regulation of immune cell function *in vivo*.

## Materials and methods

### Ethics

Animal care and use was approved by the Institutional Animal Care and Use Committee of University of Wisconsin and strictly followed guidelines set by the federal Health Research Extension Act and the Public Health Service Policy on the Humane Care and Use of Laboratory Animal, administered by the National Institute of Health Office of Laboratory Animal Welfare.

### Zebrafish husbandry

All protocols using zebrafish and mouse in this study have been approved by the University of Wisconsin-Madison Research Animals Resource Center (protocols M005405-A02/zebrafish, M005916/mouse). Adult zebrafish were maintained on a 14 h:10 h light/dark schedule. Upon fertilization, embryos were transferred into E3 medium and maintained at 28.5°C. To prevent pigment formation, larvae were maintained in E3 medium containing 0.2 mM *N*-Phenylthiourea (PTU) (Sigma-Aldrich, St. Louis, MO) starting at 1 dpf. Adult wild-type AB strain zebrafish and transgenic zebrafish lines including *Tg(tnf:GFP)*(40) and *Tg(mpeg1:Cherry-CAAX)*(47) were utilized in this study.

### Bacterial culture and preparation

Unlabeled or mCherry-expressing *Listeria monocytogenes* strain 10403 was used in this study. *L. monocytogenes* were grown in brain–heart-infusion (BHI) medium (Becton, Dickinson and Company, Sparks, MD). A streak plate from frozen stock was prepared and grown overnight at 37°C; the plate was stored at 4°C. The day before infection, a fresh colony was picked from the streak plate and grown statically in 1 mL BHI overnight at 30°C to reach stationary phase and to flagellate bacteria. The next day bacteria were prepared to infect either primary macrophages or zebrafish larvae. To prepare for infection of primary cells, the 1 mL suspension was diluted with 1 mL sterile PBS, OD was determined to calculate the number of bacteria to infect cells at multiplicity of infection (MOI) of 2 (1 cell: 2 bacteria) (OD 1= 7.5 × 10^8^ bacteria). To prepare for zebrafish tail wound infection, bacteria were sub-cultured for ∼1.5-2.5 h in fresh BHI (1:4 culture:BHI; 5 mL total) to achieve growth to mid-logarithmic phase (OD600 ≈ 0.6-0.8). From this sub-cultured bacterial suspension, 1 mL aliquot was collected, spun down at high speed for 30 seconds at room temperature, washed three times in sterile PBS and resuspended in 100 µL of sterile PBS. Unlabeled mCherry-labeled bacteria

### *L. monocytogenes* infection of mouse bone marrow derived macrophages (BMDM)

Six- to 8-week-old C57BL/6 female mice were obtained from NCI/Charles River NCI facility and bone marrow-derived macrophages were made as previously described (48). Briefly, macrophages were cultured from bone marrow in the presence of M-CSF derived from transfected 3T3 cell supernatant for 6 days, with an additional supplement of M-CSF medium 3 days postharvest. Cells were frozen down for storage. The day before infection, frozen cells were thawed and plated in 35 mm glass bottom dishes (MatTek, Ashland, MA) at 1.6 × 10^6^ in 2.4 mL BMDM medium (RPMI containing 10% fetal bovine serum, 10% CSF, 1% sodium pyruvate, 1% glutamate and 0.1% β-mercaptoethanol) and allowed to recover overnight at 37°C, 5% CO_2_. The following day 1.6 mL BMDM medium with or without *L. monocytogenes* at MOI 2 was added to cells and incubated at 37°C and 5% CO_2_. After 30 minutes, cells were rinsed once with BMDM medium and replaced with 2.4 mL medium containing 0.25 mg/mL gentamicin (Lonza, Walkersville, MD). Cells were maintained at 37°C and 5% CO_2_ and imaged live by FLIM at 5-6 hours post infection.

### Zebrafish tail wounding

Simple tail fin transection, infected tail fin transection and thermal injury of the tail fin were performed on 3 days post fertilization (dpf) larvae as described previously (38). In preparation for wounding, larvae were anesthetized in E3 medium containing 0.16 mg/mL Tricaine (ethyl 3-aminobenzoate; Sigma-Aldrich). Simple tail transection of the caudal fin was performed using surgical blade (Feather, no. 10) at the boundary of and without injuring the notochord; following transection, larvae were rinsed with E3 medium to wash away Tricaine, placed in fresh milk-coated dishes with fresh E3 medium, and maintained at 28.5°C until live imaging. For infected tail transection, larvae were placed in 5 mL E3 medium containing Tricaine in 60-mm milk-treated dish; 100 µL unlabeled bacterial suspension in PBS, or 100 µL PBS for control uninfected wounding, was added to the E3 medium and swirled gently to achieve even distribution of bacteria; tail fin transection of larvae in control or infected E3 medium was performed as described above; larvae were immediately transferred to a horizontal orbital shaker and shaken for 30 min at 70-80 rpm; control and infected larvae were then rinsed five times with 5 mL E3 medium without Tricaine to wash away bacteria and maintained at 28.5°C until live imaging; larvae were not treated with antibiotics at any point during the experiment. To perform thermal injury, fine tip (type E) of a line-powered thermal cautery instrument (Stoelting, Wood Dale, IL) was placed into the E3 medium, held to the posterior tip of the caudal fin, and turned on for 1-2 s until tail fin tissue curled up without injuring the notochord; following injury, larvae were rinsed with E3 medium to wash away Tricaine, placed in fresh milk-coated dishes with fresh E3 medium, and maintained at 28.5°C until live imaging.

### Inhibition of glycolysis in wounded zebrafish

2-deoxy-d-glucose (2-DG; Sigma-Aldrich, St. Louis, MO) treatment of wounded zebrafish larvae (simple tail transection) was empirically optimized by testing different doses and length of pretreatment. 2-DG is a weak, but fast acting inhibitor. Inhibitor was freshly prepared for each experiment, dissolved at 100 mM in E3 medium. Treatment was performed by bathing larvae in E3 medium containing 5 mM 2-DG for 1 hour before imaging. Larvae were kept in the presence of the inhibitor during imaging.

### Embedding zebrafish larvae for live imaging

Larvae were embedded in 1 mL 1% low gelling agarose prepared in E3 medium in Ibidi µ-Slide 2-Well Glass Bottom Chamber (Ibidi, Fitchburg, WI) and topped off with 1 mL E3 medium. Agarose and top-off solution were supplemented with 0.16 mg/mL Tricaine to keep larvae anesthetized during imaging. In 2-DG experiments, agarose and top-off solution were supplement with 5 mM 2-DG to maintain larvae in the inhibitor.

### Fluorescence Lifetime Imaging of NAD(P)H and FAD

All samples were imaged using a 2-photon fluorescence microscope (Ultima, Bruker) coupled to an inverted microscope body (TiE, Nikon), adapted for fluorescence lifetime acquisition with time correlated single photon counting electronics (SPC-150, Becker & Hickl, Berlin, Germany). A 40X (NA=1.15) water immersion objective was used. An Insight DS+ (Spectra Physics) femtosecond source with dual emission provided light at 750 nm (average power: 1.4 mW) for NAD(P)H excitation and 1040 nm (average power: 2.1 mW) for mCherry excitation. FAD excitation at 895 nm was achieved through wavelength mixing. Wavelength mixing was achieved by spatially and temporally overlapping two synchronized pulse trains at 750 nm and 1040 nm (36). Bandpass filters were used to isolate light, with 466/40 nm used for NAD(P)H and 540/24 nm for FAD, and 650/45 for mCherry which were then detected by GaAsP photomultiplier tubes (H7422, Hamamatsu). Fluorescence lifetime decays of NAD(P)H, FAD, and mCherry were acquired simultaneously with 256 time bins across 256×256 pixel images within Prairie View (Bruker Fluorescence Microscopy) with a pixel dwell time of 4.6 µs and an integration time of 60 seconds at an optical zoom of 2.00. No change in the photon count rate was observed, ensuring that photobleaching did not occur. The second harmonic generation obtained from urea crystals excited at 890 nm was used as the instrument response function and the full width at half maximum was measured to be 260 ps. BMDM were imaged live in MatTek dishes while maintained at 37°C and 5% CO_2_ using a stage top incubator system (Tokai Hit, Bala Cynwyd, PA). Zebrafish larvae were imaged live at room temperature.

### Fluorescence Lifetime Data Analysis

Fluorescence lifetime components were computed in SPCImage v7.4 (Becker and Hickl). For each image, a threshold was selected to exclude background. The fluorescence lifetime components were then computed for each pixel by deconvolving the measured instrument response function and fitting the resulting exponential decay to a two-component model, I(t) = α_1_e^−t/τ1^ + α_2_e^−t/τ2^ + C, where I(t) is the fluorescence intensity at time t after the laser excitation pulse, α_1_ and α_2_ are the fractional contributions of the short and long lifetime components, respectively (i.e., α_1_ + α_2_ = 1), τ_1_ and τ_2_ are the fluorescence lifetimes of the short and long lifetime components, respectively, and C accounts for background light. A two-component decay was used to represent the lifetimes of the free and bound configurations of NAD(P)H and FAD (22, 49). Images were analyzed at the single cell level. For the *in vitro* macrophages, cell cytoplasm masks were obtained using a custom CellProfiler pipeline (v.3.1.8) (50). Briefly, the user manually outlined the nucleus of the cells and those masks were then propagated outwards to find cell areas. Cytoplasm masks were then determined by subtracting the nucleus masks from the total cell area masks. Bacteria masks were created in Fiji (51) by thresholding the mCherry intensity images into bacteria and background. The resulting bacteria masks were then subtracted from the corresponding field of view’s masks to exclude bacterial metabolic data. The diffuse cytoplasmic fluorescence in the mCherry images is likely due to FAD autofluorescence (44). Images of the optical redox ratio (intensity of NAD(P)H divided by the sum of the intensity of NAD(P)H and the intensity of FAD) and the mean fluorescence lifetime (τ_m_ = α_1_τ_1_ + α_2_τ_2_, where τ_1_ is the short lifetime for free NAD(P)H and bound FAD, τ_2_ is the long lifetime of bound NAD(P)H and free FAD, and α_1_ and α_2_ represent relative contributions from free and protein-bound NAD(P)H respectively, and the converse for FAD α_1_ and α_2_) of NAD(P)H and FAD were calculated and autofluorescence imaging endpoints were averaged for all pixels within a cell cytoplasm using RStudio v. 1.2.1335 (52). For the *in vivo* macrophages, a custom CellProfiler pipeline segmented the macrophage cell area. Briefly, the pipeline rescaled the mCherry intensity images to be between 0 and 1 by dividing by the brightest pixel value in the image. Background was excluded by manually setting a threshold (0.15). Cells were identified using CellProfiler’s default object identification. Then, each cell was manually checked and edited as necessary to exclude background fluorescence and to include all pixels of each macrophage. Images of the optical redox ratio (intensity of NAD(P)H divided by the sum of the intensity of NAD(P)H and the intensity of FAD) and the mean fluorescence lifetime (τ_m_ = α_1_τ_1_ + α_2_τ_2_; defined above) of NAD(P)H and FAD were calculated and autofluorescence imaging endpoints were averaged for all pixels within a cell using MATLAB v.9.7.01296695 (R2019b; Mathworks, Natick, MA).

### Metabolomics

To analyze intracellular metabolites, metabolites were extracted with cold liquid chromatography–mass spectrometry (LC–MS) grade 80/20 methanol/H2O (v/v). Samples were dried under nitrogen flow and subsequently dissolved in LC–MS grade water for LC–MS analysis methods. Protein pellets were removed by centrifugation. Samples were analyzed using a Thermo Q-Exactive mass spectrometer coupled to a Vanquish Horizon Ultra-High Performance Liquid Chromatograph (UHPLC). Metabolites were separated on a C18 (details below) at a 0.2 ml per min flow rate and 30 °C column temperature. Data was collected on full scan mode at a resolution of 70 K. Samples were loaded in water and separated on a 2.1 × 100 mm, 1.7 μM Acquity UPLC BEH C18 Column (Waters) with a gradient of solvent A (97/3 H2O/methanol, 10 mM TBA, 9 mM acetate, pH 8.2) and solvent B (100% methanol). The gradient was: 0 min, 5% B; 2.5 min, 5% B; 17 min, 95% B; 21 min, 95% B; 21.5 min, 5% B. Data were collected on a full scan negative mode. The identification of metabolites reported was based on exact m/z and retention times, which were determined with chemical standards. Data were analyzed with Maven. Relative metabolite levels were normalized to protein content.

### Statistical analyses

Biological repeats are defined as separate clutches of embryos collected on separate days. Statistical analyses were performed using R v.3.6.2 (www.R-project.org)(53). General linear models were fit to data, where every data point represented a macrophage. Models included *day* (biological repeat) as a blocking factor. An interaction was included in all models where more than one experimental factor was present (for example, time and treatment), to determine whether effects associated with experimental factors modified one another. All models utilized cluster-robust standard errors to account for multiple macrophages being measured within the same larvae. Log transformation was applied to certain lifetime endpoints prior to analysis to improve scaling and symmetry, and lessen the influence of outliers. No adjustment for multiplicity was done. Statistical significance was set to 0.05. All data were graphed using Prism (GraphPad Software, Inc., San Diego, CA). Each data point in the graphical display represent a macrophage and the data for each condition is presented as a composite dotplot and boxplot; each biological repeat is displayed by a different color in the dotplot; boxplots show median (central line), first and third quartiles (lower and upper lines), and the Tukey method was employed to create the whiskers (the farthest data points that are no further than 1.5 times the interquartile range); data points beyond whiskers (refer to dotplot) are considered outliers. Graphical information involving zebrafish experiments is accompanied by statistical conclusions (below graphs) that account for the cluster-correlated structure of data as described above.

## Data and code availability

Data generated and analyzed, including codes and algorithms related to the analysis of data in the current study are available from the corresponding authors upon request.

## Acknowledgements

We would like to thank members of the Huttenlocher and Skala laboratories, notably Jayne M Squirrell, Elizabeth Berge, Steve Trier, Tiffany M Heaster and Amani Gillette, for valuable discussions and technical assistance over the course of this work.

## Grant Support

This work is supported by R35 GM118027 to AH, R01 CA205101 to MCS, and individual fellowship from American Heart Association to VM (17POST33410970).

## Figure supplement legends

**Figure 1 supplement.**
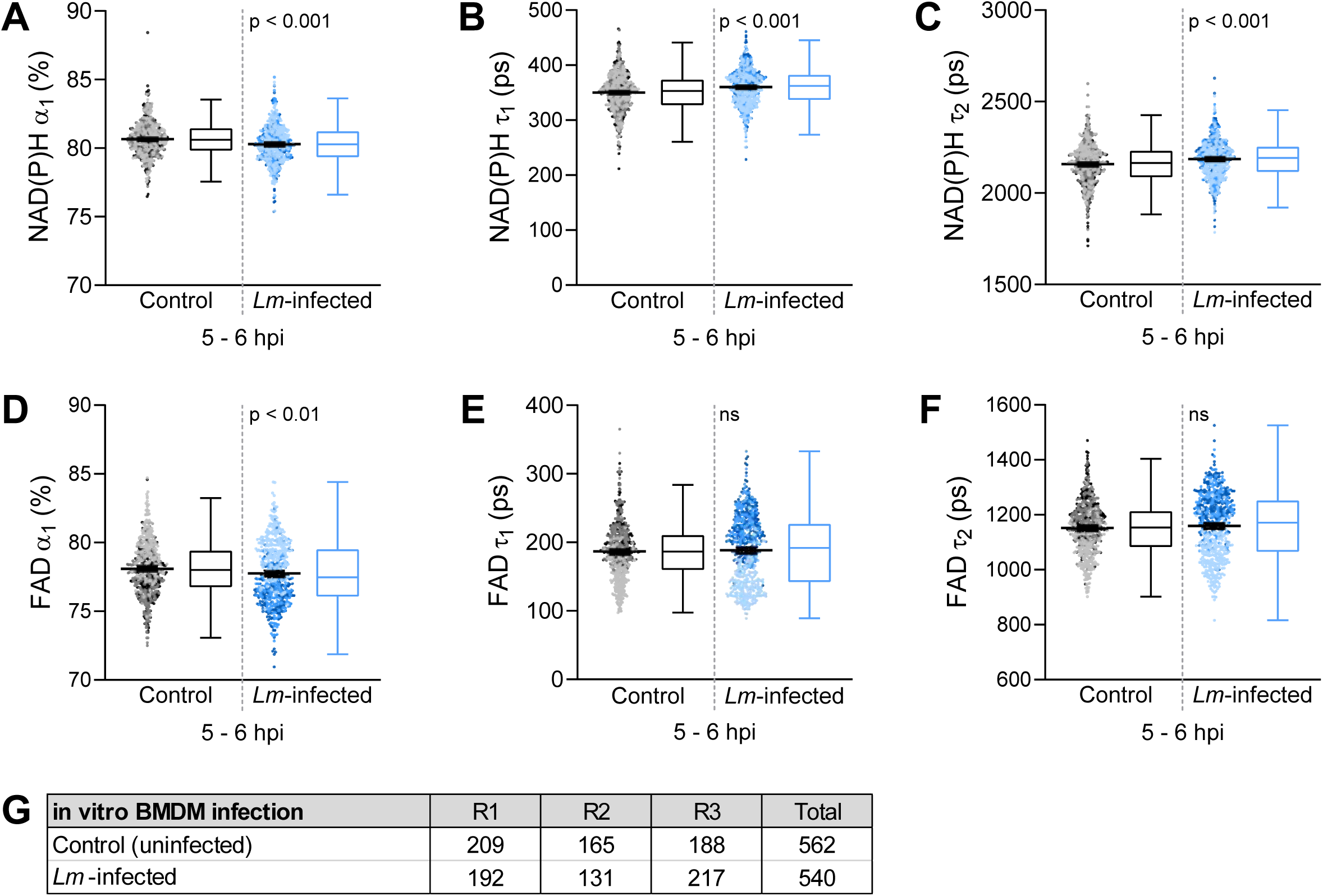
Other NAD(P)H and FAD fluorescence lifetime components measured during *in vitro Listeria monocytogenes (Lm)* infection of primary mouse macrophages, associated with Figure 1. Quantitative analysis of A) alpha1 (α_1_), fractional component of free NAD(P)H; B) tau1 (τ_1_), free/short lifetime of NAD(P)H; C) tau2 (τ_2_), bound/long lifetime of NAD(P)H; D) alpha1 (α_1_), fractional component of bound FAD; E) tau1 (τ_1_), bound/short lifetime of FAD; F) tau2 (τ_2_), free/long lifetime of FAD. Each data point represent a macrophage and the data for each condition is displayed by a composite dotplot and boxplot; each repeat (n=3) is displayed by a different color in the dotplot, showing mean with 95% CI; boxplots show median (central line), first and third quartiles (lower and upper lines), and the Tukey method was employed to create the whiskers; data points beyond whiskers (refer to dotplot) are considered outliers. Statistical comparison was performed by general linear model; ns = not significant. G) Sample size of data set shown in Figure 1 and this supplement.

**Figure 2 supplement.**
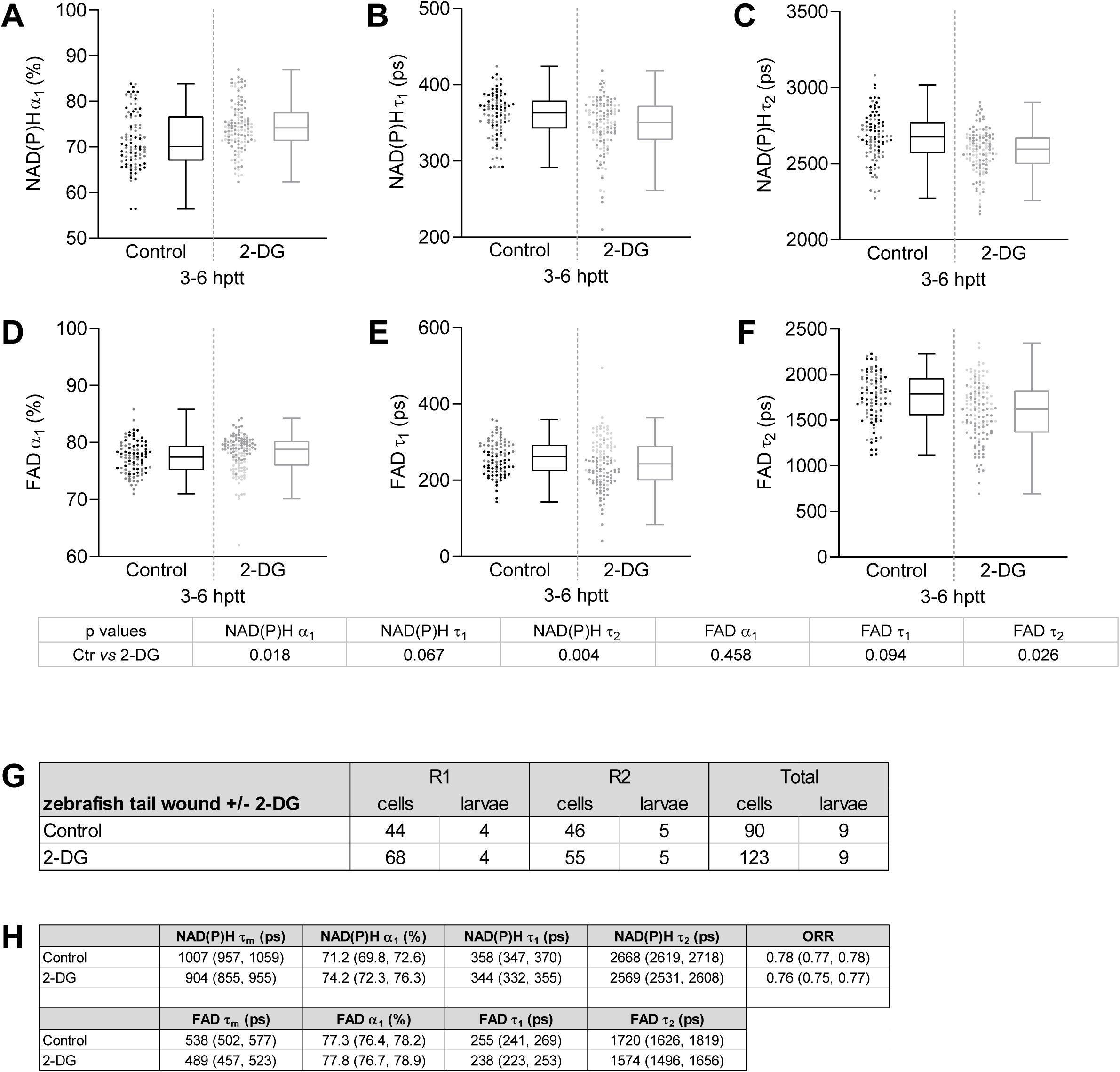
Other NAD(P)H and FAD fluorescence lifetime components measured in macrophages at simple tail wounds of zebrafish larvae treated with glycolysis inhibitor (2-DG), associated with Figure 2. Quantitative analysis of A) alpha1 (α_1_), fractional component of free NAD(P)H; B) tau1 (τ_1_), free/short lifetime of NAD(P)H; C) tau2 (τ_2_), bound/long lifetime of NAD(P)H; D) alpha1 (α_1_), fractional component of bound FAD; E) tau1 (τ_1_), bound/short lifetime of FAD; F) tau2 (τ_2_), free/long lifetime of FAD. Each data point represent a macrophage and the data for each condition is displayed by a composite dotplot and boxplot; each repeat (n=2) is displayed by a different color in the dotplot; boxplots show median (central line), first and third quartiles (lower and upper lines), and the Tukey method was employed to create the whiskers; data points beyond whiskers (refer to dotplot) are considered outliers. Statistical comparison was performed using a general linear model with cluster-robust standard errors to account for multiple macrophages measured per larvae, thereby statistical conclusions are shown in a table below the graphs. All lifetime endpoints were log-transformed prior to analysis. G) Sample size of data set shown in Figure 2 and this supplement. H) Estimated means with 95% CI; ORR = optical redox ratio.

**Figure 3 supplement.**
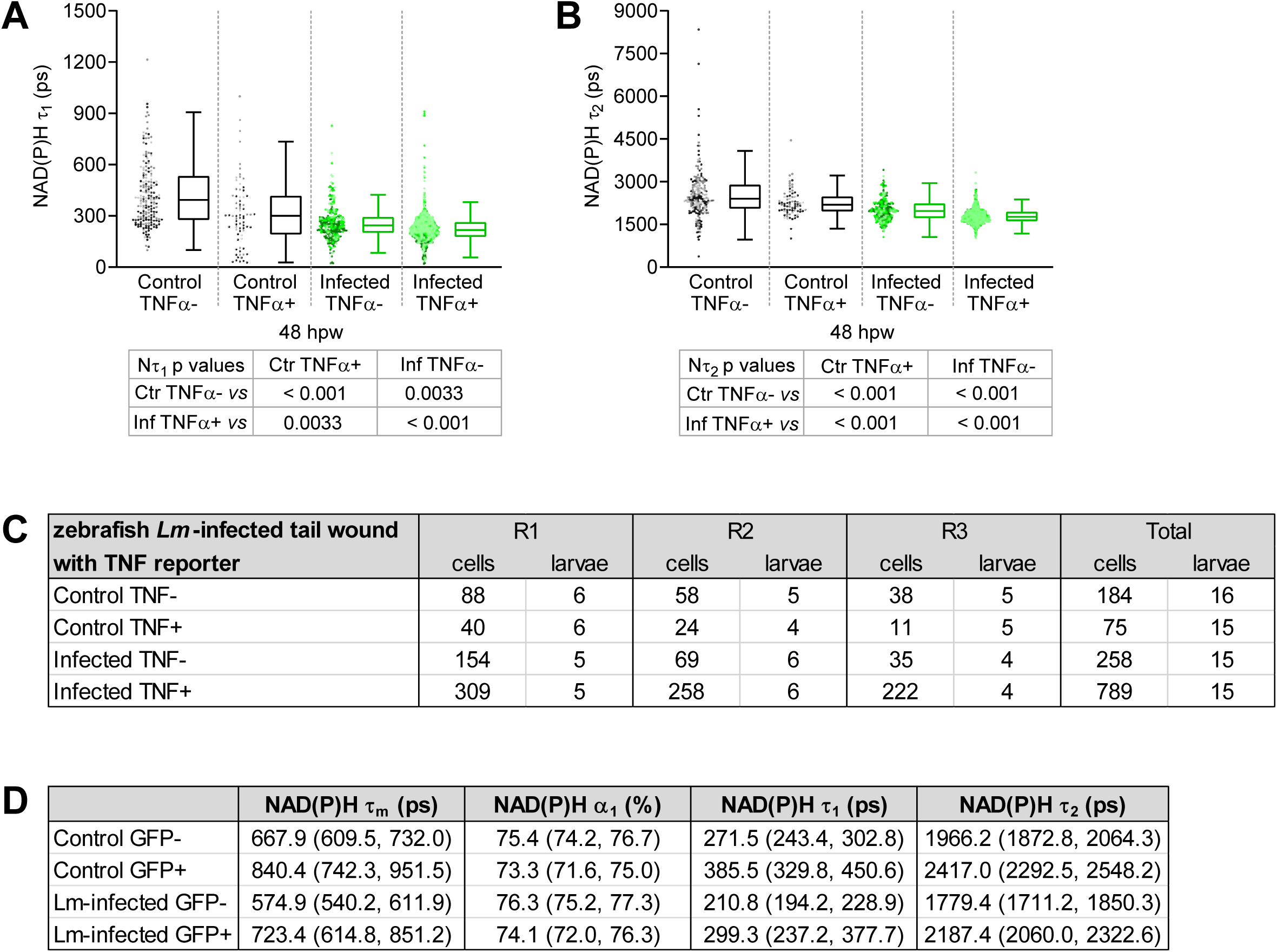
Other NAD(P)H fluorescence lifetime components measured in TNFα− and TNFα+ macrophages at infected tail wounds of zebrafish larvae, associated with Figure 3. Quantitative analysis of A) tau1 (τ_1_), free/short lifetime of NAD(P)H; B) tau2 (τ_2_), bound/long lifetime of NAD(P)H. Each data point represent a macrophage and the data for each condition is displayed by a composite dotplot and boxplot; each repeat (n=3) is displayed by a different color in the dotplot; boxplots show median (central line), first and third quartiles (lower and upper lines), and the Tukey method was employed to create the whiskers; data points beyond whiskers (refer to dotplot) are considered outliers. Statistical comparison was performed using a general linear model with cluster-robust standard errors to account for multiple macrophages measured per larvae, thereby statistical conclusions are shown in a table below the graphs. The lifetime endpoints were log-transformed prior to analysis. Interaction between treatment and GFP expression was included to analyze whether either factor modified the effect of the other; no interaction was found. C) Sample size of data set shown in Figure 3 and this supplement. D) Estimated means with 95% CI.

**Figure 4 supplement.**
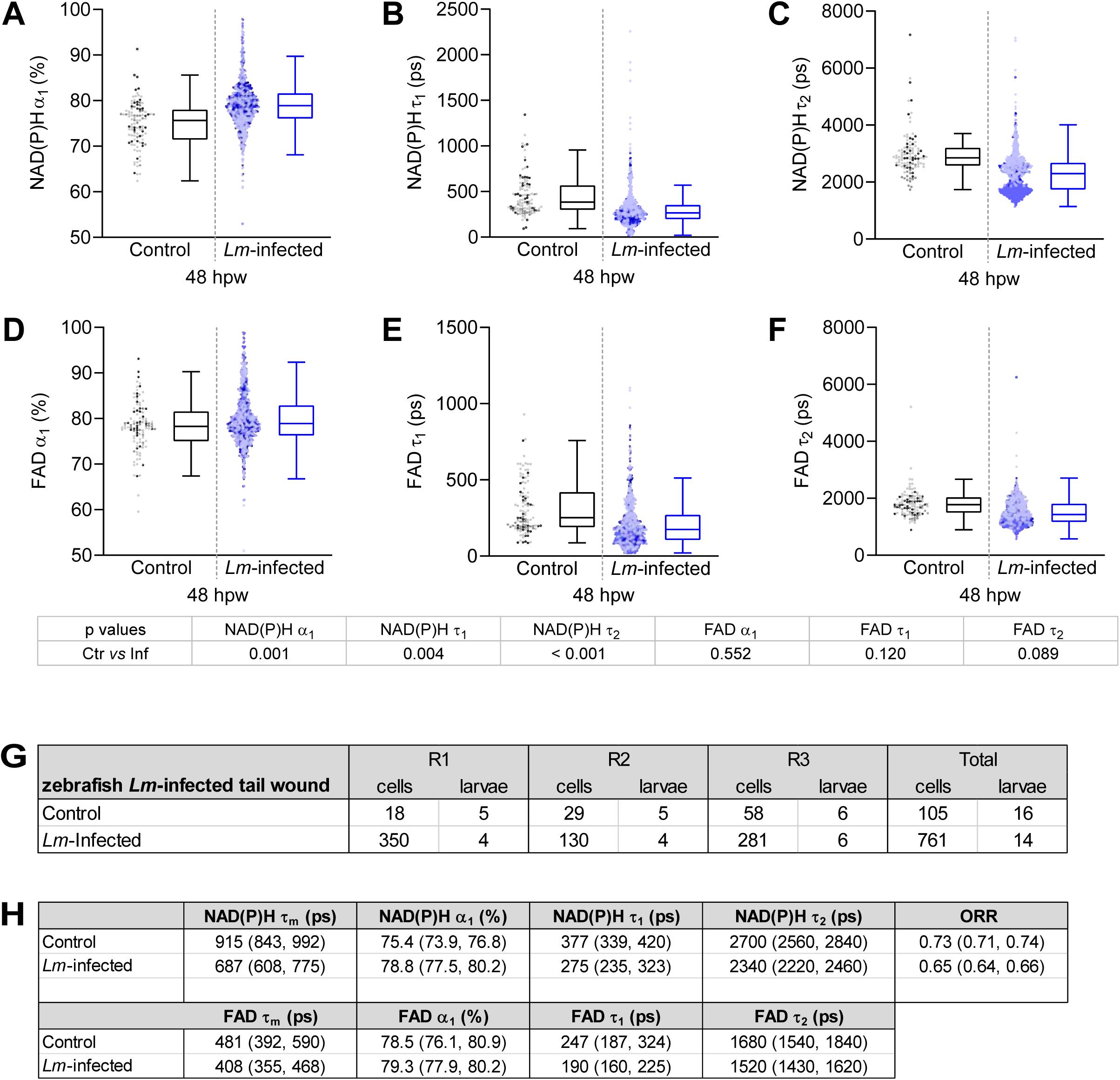
Other NAD(P)H and FAD fluorescence lifetime components measured in macrophages at infected tail wounds of zebrafish larvae, associated with Figure 4. Quantitative analysis of A) alpha1 (α_1_), fractional component of free NAD(P)H; B) tau1 (τ_1_), free/short lifetime of NAD(P)H; C) tau2 (τ_2_), bound/long lifetime of NAD(P)H; D) alpha1 (α_1_), fractional component of bound FAD; E) tau1 (τ_1_), bound/short lifetime of FAD; F) tau2 (τ_2_), free/long lifetime of FAD. Each data point represent a macrophage and the data for each condition is displayed by a composite dotplot and boxplot; each repeat (n=3) is displayed by a different color in the dotplot; boxplots show median (central line), first and third quartiles (lower and upper lines), and the Tukey method was employed to create the whiskers; data points beyond whiskers (refer to dotplot) are considered outliers. Statistical comparison was performed using a general linear model with cluster-robust standard errors to account for multiple macrophages measured per larvae, thereby statistical conclusions are shown in a table below the graphs. Log transformation was applied to τ_1_ and τ_2_ prior to analysis. G) Sample size of data set shown in Figure 4 and this supplement. H) Estimated means with 95% CI; ORR = optical redox ratio.

**Figure 5 supplement.**
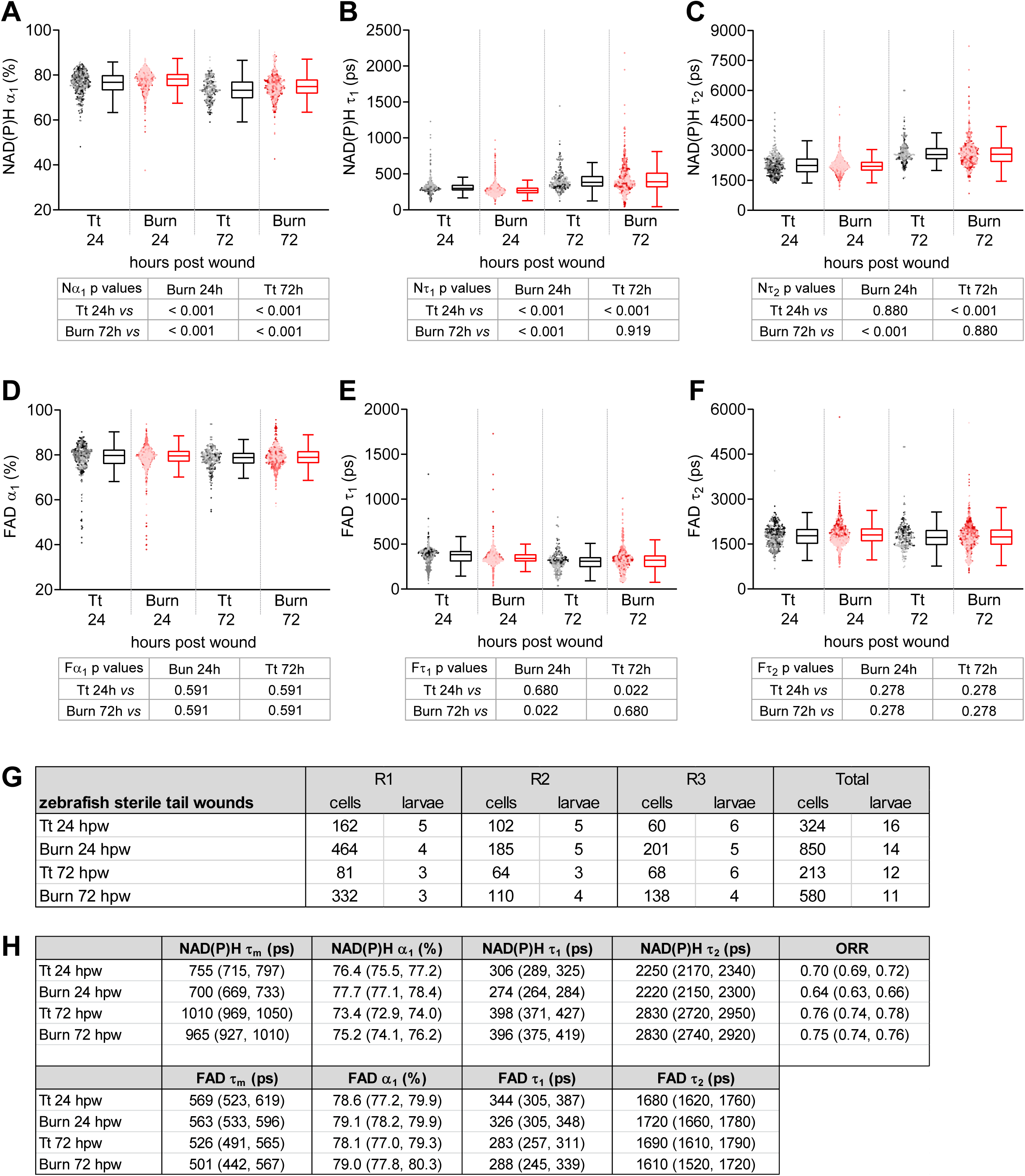
Other NAD(P)H and FAD fluorescence lifetime components measured in macrophages at sterile tail wounds of zebrafish larvae, associated with Figure 5. Quantitative analysis of A) alpha1 (α_1_), fractional component of free NAD(P)H; B) tau1 (τ_1_), free/short lifetime of NAD(P)H; C) tau2 (τ_2_), bound/long lifetime of NAD(P)H; D) alpha1 (α_1_), fractional component of bound FAD; E) tau1 (τ_1_), bound/short lifetime of FAD; F) tau2 (τ_2_), free/long lifetime of FAD. Each data point represent a macrophage and the data for each condition is displayed by a composite dotplot and boxplot; each repeat (n=3) is displayed by a different color in the dotplot; boxplots show median (central line), first and third quartiles (lower and upper lines), and the Tukey method was employed to create the whiskers; data points beyond whiskers (refer to dotplot) are considered outliers. Statistical comparison was performed using a general linear model with cluster-robust standard errors to account for multiple macrophages measured per larvae, thereby statistical conclusions are shown in a table below the graphs. Log transformation was applied to τ_1_ and τ_2_ prior to analysis. Interaction between treatment and time was included to analyze whether either factor modified the effect of the other; weak interaction was detected for NAD(P)H τ_1_. G) Sample size of data set shown in Figure 5 and this supplement. H) Estimated means with 95% CI; ORR = optical redox ratio.

## Notes

### Competing Interest Statement

The authors have declared no competing interest.

